# In vivo inhibition of stearoyl-CoA desaturase modulates the hippocampal fatty acid profile and restores density of dendritic spines in the aggressive 5xFAD model of Alzheimer’s disease

**DOI:** 10.1101/2025.08.27.672447

**Authors:** Marta Turri, Myriam Aubin, Anne Aumont, Annick Vachon, Jessica Avila Lopez, Marian Mayhue, Mélanie Plourde, Karl Fernandes

## Abstract

While alterations in brain lipids are a central feature of Alzheimer’s disease (AD), therapeutic strategies targeting brain lipid metabolism are still lacking. Prior preclinical work has shown that pharmacological inhibition of the fatty acid desaturase, stearoyl-CoA desaturase (SCD), leads to recovery of hippocampal synapses with associated improvements in learning and memory in the slow-progressing 3xTg AD mouse model. Here, we used the rapidly progressing, highly amyloidogenic 5xFAD AD model to further delve into the effect of the SCD inhibitor (SCDi) on AD-associated fatty acid alterations and synapse loss. Hippocampus, cortex and plasma samples were collected from male and female 5xFAD and non-carrier control mice for fatty acid profiling and assessment of disease hallmarks. Plaque pathology, gliosis, and fatty acid alterations that included an increase in the C16:1/C16:0 desaturation index, a measure of SCD enzymatic activity, were apparent in the female hippocampus at 5 months of age, with similar fatty acid changes appearing in males by 8 months. Intracerebroventricular infusion of SCDi via osmotic pump for 28 days in 5 months old female 5xFAD and NC mice modulated the SCD-related fatty acid disturbances as well as PUFA concentrations. Quantification of Golgi staining revealed an SCDi-induced recovery of dendritic spine density. The beneficial effects of SCDi treatment on fatty acid balance and hippocampal dendritic spines in this more aggressive amyloidogenic 5xFAD model further support SCD inhibition as a promising therapeutic avenue for AD.

## Introduction

Alzheimer’s disease (AD) is a complex and multifactorial neurodegenerative disorder that is characterized by progressive brain dysfunctions that lead to incurable cognitive decline. Familial AD (FAD), which is an aggressive and relatively rare early-onset form, is caused by autosomal dominant mutations in the genes for amyloid precursor protein (APP), Presenilin-1 (PSEN1) or PSEN2, and is associated with altered processing of ß-amyloid peptides. Sporadic AD (SAD), which tends to be later onset and accounts for most cases, is virtually indistinguishable from FAD pathologically and is associated with dozens of risk-enhancing genetic polymorphisms. Beyond its genetic components, AD is a multifactorial disease whose development is strongly influenced by age-related changes, vascular disease and mitochondrial function^1^. Lipid metabolism alterations are prominent features in AD and have been linked to core AD processes such as APP processing, β-amyloid deposition and clearance, tau phosphorylation and innate immunity^2–5^. Moreover, many of the major SAD risk genes are also lipid- and immunity-related. For instance, risk for SAD is increased 4-20-fold by the apolipoprotein E4 (apoE4) polymorphism of the APOE gene, implicating a lipid transport protein as the single strongest genetic risk determinant of SAD^6–8^. More generally, genome-wide meta-analyses strongly implicate lipid metabolism genes and pathways in influencing AD risk^9–11^, and AD risk genes influence blood lipid markers in patients^12^. Apart from the genetic lipid risk factors for AD, lipid-modulating metabolic and inflammatory conditions such as mid-life obesity, insulin resistance, hyperglycemia and diabetes can also increase AD susceptibility^13–17^. Despite the extensive evidence implicating lipid dysregulation in AD pathogenesis, modulating lipid metabolism remains largely undeveloped as a therapeutic strategy. This is due in part to the challenge of dissecting the contributions of the tens of thousands of distinct species of biological lipids and the diverse cellular pathways they are involved in.

One lipid pathway of emerging interest in AD relates to fatty acid metabolism. Studies on large AD cohorts have highlighted imbalances in plasma fatty acids that include increased monounsaturated fatty acids (MUFA) in patients^18^. The rate-limiting enzyme in the conversion of saturated fatty acids to MUFA, stearoyl-CoA desaturase (SCD), exhibits increased activity and expression in brains of AD patients and is associated with disease progression and decreased cognitive performance^19–22^. In past work, we identified MUFA-rich triglycerides (TG) accumulating in the periventricular zone of both post-mortem AD human brains and in the 3xTg mouse model of AD. In the 3xTg-AD mouse model, these MUFA-rich TG could be detected in advance of amyloid plaques, neurofibrillary tangles, or cognitive defect^23^. Infusion of an SCD inhibitor (SCDi) into the ventricles of pre-symptomatic AD mice helped normalize MUFA levels and led to rescue of a variety of pathological features including impaired neural stem cell activity, dendritic spine loss, microglial alterations, and deficits in spatial learning and memory^23,24^. Given the fundamental roles of fatty acids as signaling molecules, energy substrates, and building blocks of complex structural lipids, an imbalance among fatty acid species might have widespread effects on cellular function, potentially explaining why modulation of this pathway impacts multiple cell types and cellular functions. Interestingly, SCD inhibition is showing beneficial effects in several models of neurodegenerative disease^24–27^.

In the present study, we investigated fatty acid imbalance and the effects of SCD inhibition in the highly aggressive 5xFAD model of AD. The previously used 3xTg-AD model, a slow developing model that only begins to exhibit amyloid plaque deposits around 12 months of age, facilitated analysis of SCD inhibition at early stages AD pathology. In contrast, the hippocampus of 5xFAD mice already shows extensive amyloid plaque deposition by 3 months of age, making this more aggressive model useful for studying the potential of SCD inhibition at a late stage of AD amyloid pathology^28^. Using 5xFAD mice, we characterized sex-specific development of fatty acid imbalances, investigated whether SCD inhibition in vivo can correct these imbalances, and assessed the impact on density of hippocampal dendritic spines (considered a close anatomical correlate of changes in cognitive function). Our findings support investigating SCD inhibition as a beneficial therapeutic strategy even at more advanced stages of AD.

## Results

### Altered hippocampal fatty acid profile in symptomatic 5xFAD mice

The aggressive and rapid progression of AD pathologies in 5xFAD mice is reported to include extensive hippocampal amyloid plaque deposits and impaired hippocampal-dependent cognitive function by 5 months of age^29^. Since immune activation and its impact on pathology development can be affected by animal housing conditions, we first characterized key pathologies within our own breeding colony. 5xFAD and non-carrier (NC) littermates were characterized by immunohistochemistry at 2, 5 and 8 months old (MO) using 3,3’-diaminobenzidine (DAB) staining for Aβ37-42 (Aβ plaques), Iba-1 (hypertrophic microglia) and GFAP (astrocytes). Aβ37-42 staining, which was predominantly intraneuronal at 2 MO, revealed extensive extracellular plaque formation in the hippocampal dentate gyrus (DG) and cortex at the 5 MO timepoint, and this remained stable in number and density at 8 MO (Supplementary Figure S1). Evidence of microgliosis likewise emerged in the cortex and DG at 5 MO, with hypertrophic microglia displaying an inflated soma and shorter, thicker ramifications (Supplementary Figure S1). Astrocytes showed no obvious difference in morphology between 5xFAD and NC mice or between timepoints in 5xFAD brains (Supplementary Figure S1). Altogether, these observations are in line with previous reports^28^.

To focus on fatty acid changes in the context of established amyloid pathology, hippocampi were collected from male and female 5xFAD and NC counterparts at 5 and 8 MO and processed for quantitative fatty acid profiling by gas chromatography with flame ionization detector (GC-FID). At 5 MO, 5xFAD females exhibited higher concentrations of C16:0 (palmitic acid), C16:1 (palmitoleic acid), C20:4 (arachidonic acid), total omega-6, total saturated fatty acids (SFA), and total polyunsaturated fatty acids (PUFA) (Figure 1 and Supplementary Table S2) versus NC. Notably, palmitic and palmitoleic acids, which are respectively a substrate and product of SCD, were both significantly higher in these 5 MO 5xFAD females versus NC females. The C16:1/C16:0 desaturation index (DI), a measure of SCD activity^30^, was likewise higher in 5 MO 5xFAD females, with the increased C16:1 and C16 DI being maintained in 8 MO females and reaching statistical significance in males by 8 MO (Supplementary Figure S2, Supplementary Tables S1 and S2). C18 fatty acids were not increased in either females or males at 5 MO or 8 MO (Figure 1, Supplementary Figure 2, Supplementary Tables S1 and S2). Thus, the fatty acid profiles of the 5xFAD hippocampus are altered by 5 MO in females and by 8 MO in males.

**Figure 1.**
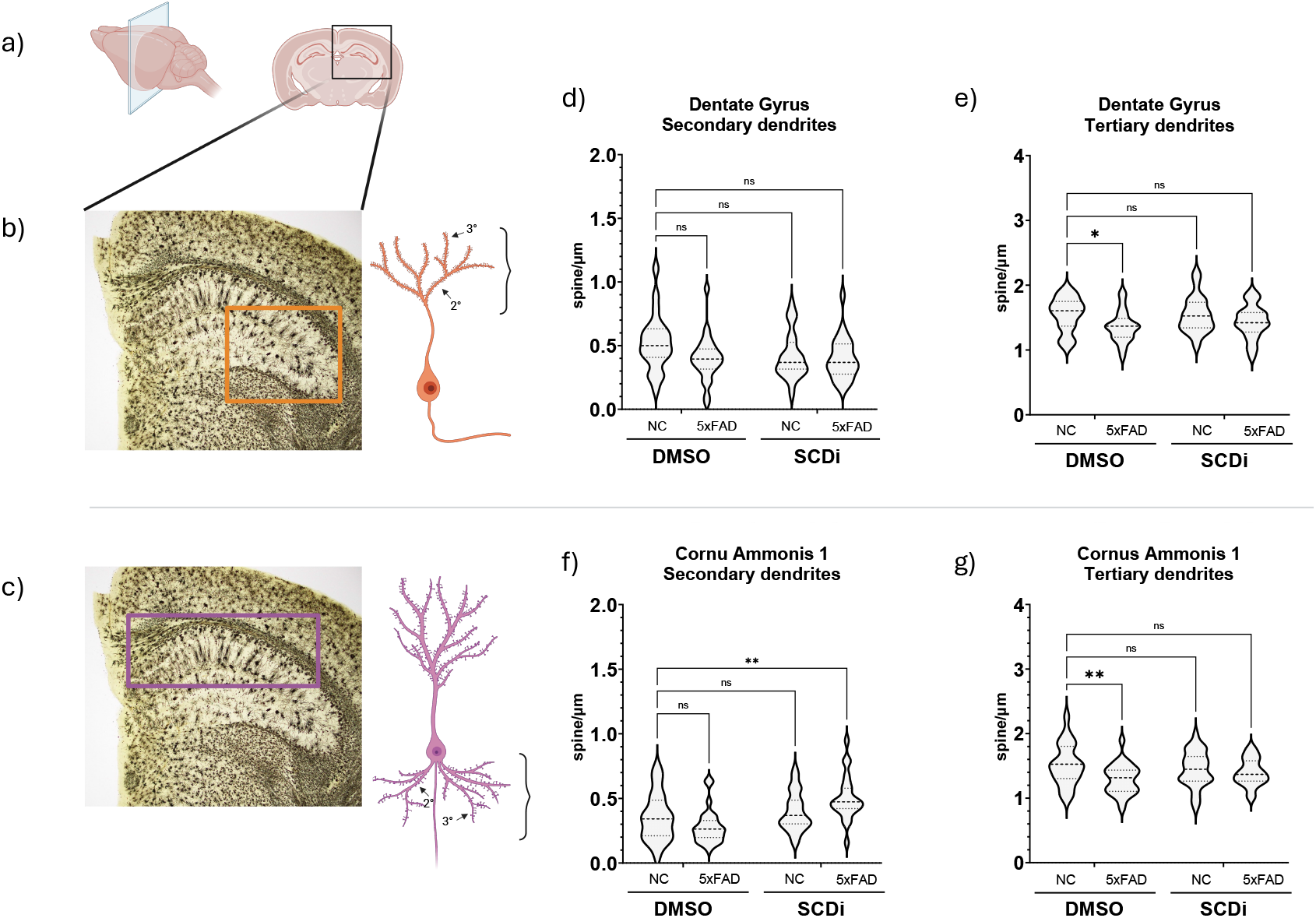
Hippocampus total fatty acid composition of 5xFAD. Column graph represents the specific fatty acid amount in hippocampus of 5xFAD (black columns, N=4) and NC (white columns, N=4) at 5 months of age. The most relevant hippocampus fatty acids are shown in male and female mice. Desaturation indexes (DI) of palmitoleic over palmitic acid, and oleic over stearic are displayed. Fatty acids are stratified by saturation degree (SFA, MUFA and PUFA) and omega-6 and omega-6/omega-3 ratio are shown. Data are expressed as mean±SEM; 5xFAD vs NC were compared by two-way ANOVA with Šídák’s multiple comparisons test as appropriate (*p<0.05 **p<0.01). C16:0, palmitic acid; C16:1, palmitoleic acid; C18:0, stearic acid; C18:1, oleic acid; C18:2, linoleic acid; C20:4, arachidonic acid, C22:6, docosahexaenoic acid; SFA, saturated fatty acids; MUFA, monounsaturated fatty acids; PUFA, polyunsaturated fatty acids.

To determine whether these fatty acid changes altered the fatty acid composition of complex lipid classes, we first separated hippocampal lipids into phospholipid, cholesteryl esters, triglycerides and free fatty acid fractions before individual GC-FID analysis. Phospholipids, which represent most brain lipids, indeed exhibited higher C16:1 acid content in females and the C16:1/16:0 DI of phospholipids was increased in both sexes (Figure 2). Cholesterol, which represents about 30% of brain lipids^31^ and can be esterified to fatty acids to form cholesteryl esters, showed similar but non-significant trends in oleic acid (C18:1) and C18 DI in females (Supplementary Figure 3). These SCD-related FA differences did not emerge in the triglycerides fraction (Supplementary Figure 3), which constitute a relatively small percentage of the brain’s overall composition, though they play an important role in brain function and metabolism^32^. Notably, the increased arachidonic acid level observed in 5xFAD females was only detectable in the cholesterol fraction (Figure Supplemental S3). Thus, specific fatty acids appear to be targeted towards particular lipid classes.

**Figure 2.**
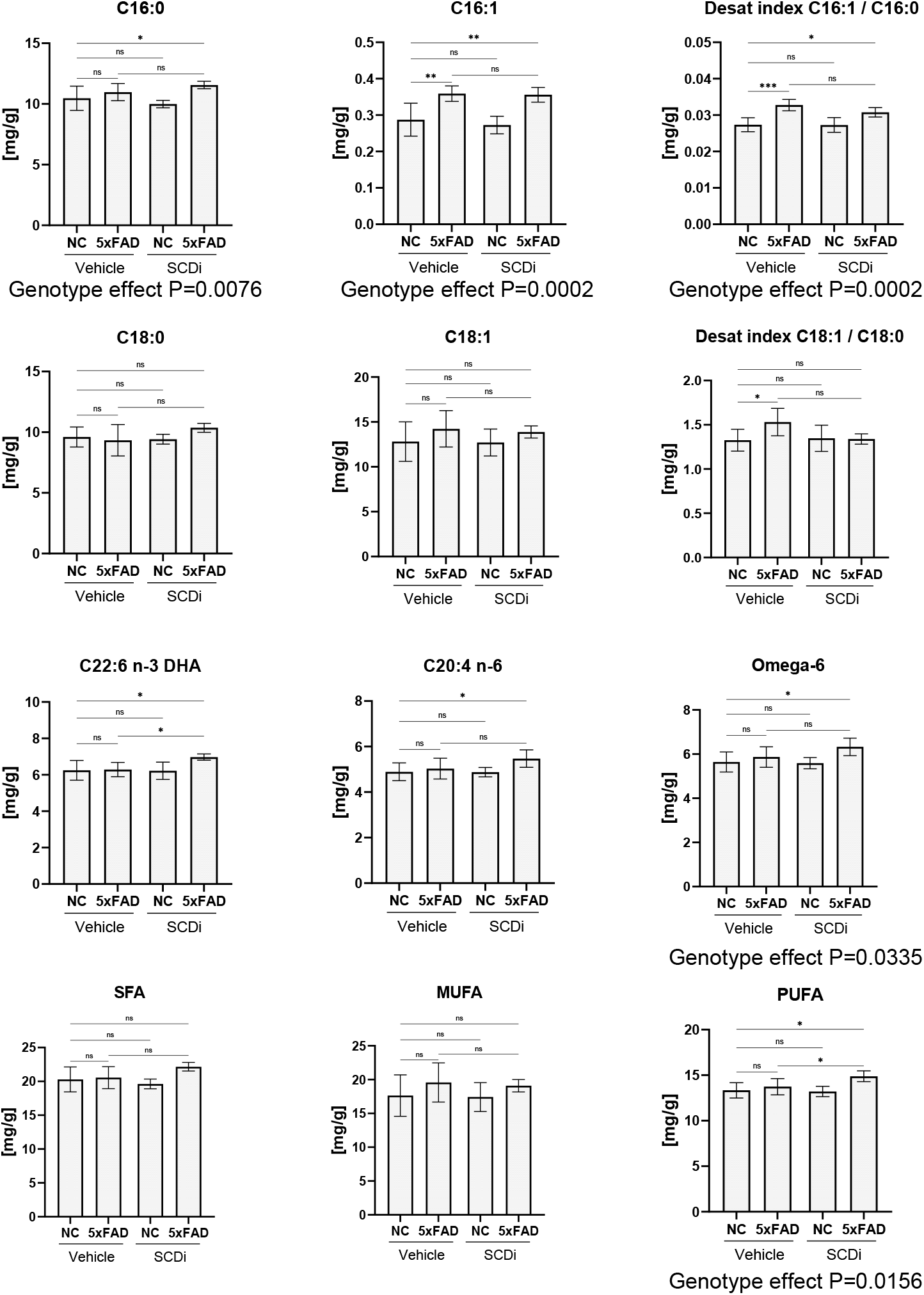
Hippocampus SCD-related phospholipids fatty acid composition of 5xFAD. Column graph represents the SCDi related fatty acid amount in phospholipids (PL) of hippocampus of 5xFAD (black columns, N=4) and NC (white columns, N=4) at 5 months of age. Desaturation indexes (DI) of palmitoleic over palmitic acid, and oleic over stearic are displayed. Data are expressed as mean±SEM; 5xFAD vs NC were compared by two-way ANOVA with Šídák’s multiple comparisons test as appropriate (*p<0.05 ***p<0.001). C16:0, palmitic acid; C16:1, palmitoleic acid; C18:0, stearic acid; C18:1, oleic acid; acid.

We also analyzed FAs in the plasma and cortex of 5xFAD and NC mice. For plasma FAs, no differences between NC and 5xFAD were present for either males or females at 2, 5, and 8 months of age (Supplementary Table S3 and S4). Age-dependent changes were present in the major brain-associated PUFAs only in 5xFAD females: AA and DHA were significantly decreased from 5 MO, while in males AA and DHA levels were higher than in females and remained relatively stable in both NC and 5xFAD across ages. We then analyzed the FA composition of the cortex at later symptomatic stages (8 MO mice). Of the 45 FAs that can be detected with the GC-FID, none were significantly altered in the cortex of mice at this age (Supplementary Table S5).

Together, these lipidomic analyses reveal sex- and age-dependent changes in hippocampal fatty acid profiles of 5xFAD mice, including in SCD-associated fatty acids and PUFA.

### Brain SCD inhibition modifies the hippocampal fatty acid profile of 5xFAD mice

Pharmacological inhibition of brain SCD activity in the slow developing 3xTg-AD model of AD neuropathology was previously reported to modulate the brain fatty acid profile and resulted in structural and functional improvements in the hippocampus^24^. Since the above GC-FID analyses of 5xFAD mice revealed fatty acid alterations, including in SCD-related fatty acids, we used a similar approach to test the impact of SCD inhibition on the fatty acid imbalance in this more aggressive neuropathology model. A commercially available SCD inhibitor (Abcam, cat# ab142089) was infused into the lateral ventricles of 5 MO female 5xFAD and NC mice for 28 days using intracerebroventricular osmotic pumps, as previously^24^, with vehicle-infused mice allowing for control of the effects of pump implantation. Vehicle-infused 5xFAD mice showed the expected higher C16:1 and C16 DI versus NC mice, with the C18 DI also reaching significance (Figure 3 and Supplementary Table S6). In the SCDi-infused mice, although C16 DI remained increased in 5xFAD mice, SCDi increased the C16:0 substrate, eliminated the increase in the C18 DI and increased total SFA, changes consistent with reduced SCD activity. Interestingly, SCDi infusion in 5xFAD mice also increased the hippocampal omega-6 AA, omega-3 DHA and total PUFA in the hippocampus of 5xFAD mice when compared to NC-SCDi (Figure 3 and Supplementary Table S6), suggesting selective effects of SCDi in the 5xFAD mice.

**Figure 3.**
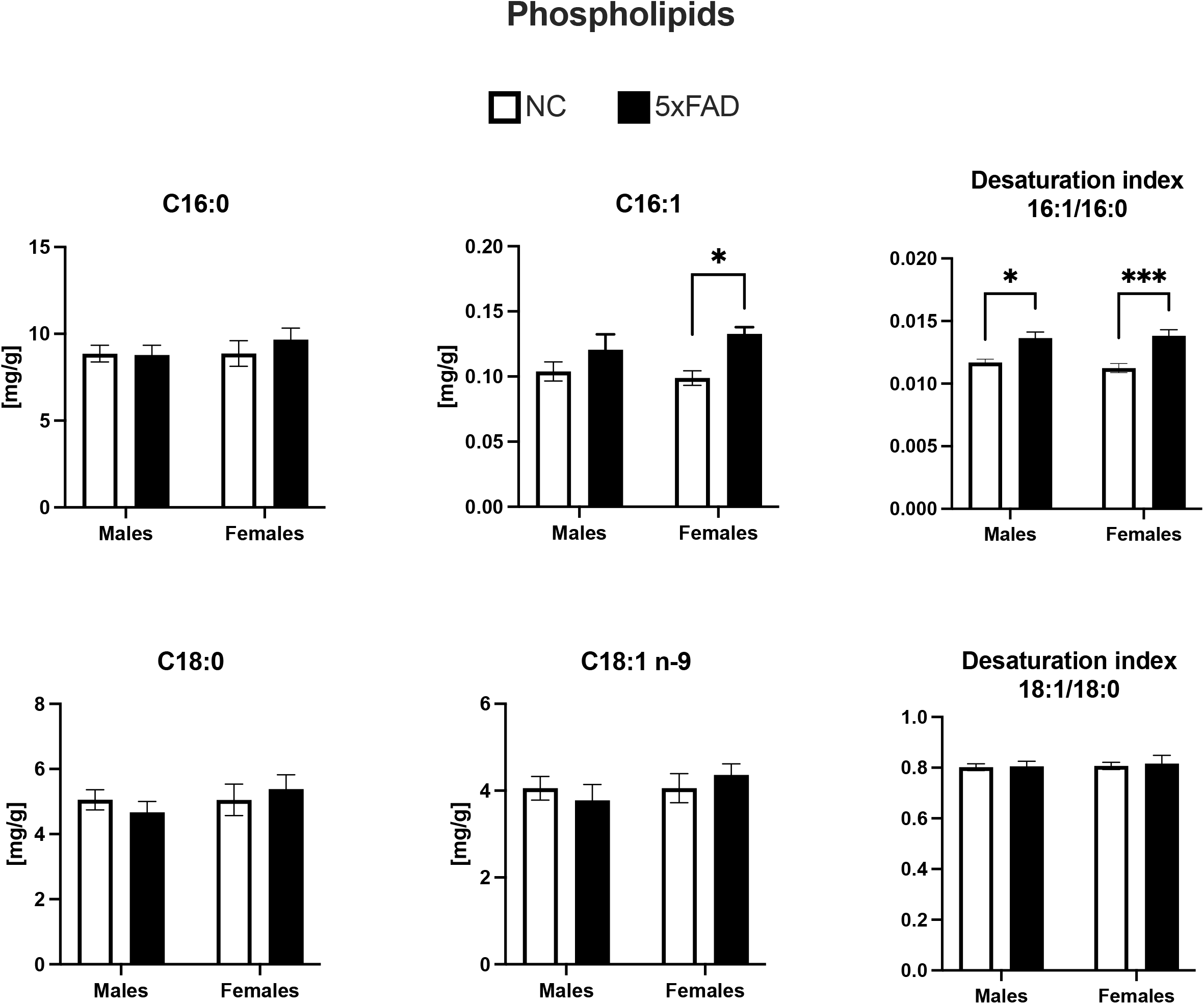
Effects of SCD inhibition on fatty acids composition of hippocampus of 5xFAD. Column graph represents the specific fatty acid amount of hippocampus of 5xFAD and NC at 6 months of age after a month of treatment. The most relevant hippocampus fatty acids are shown in SCDi and Vehicle-treated mice (4 mice per group: N=4 NC-Vehicle, N=4 NC-SCDi, N=4 5xFAD-Vehicle, N=4 5xFAD-SCDi). Desaturation indexes (DI) of palmitoleic over palmitic acid, and oleic over stearic acid are displayed. Fatty acids are stratified by saturation degree (SFA, MUFA and PUFA). Data are expressed as mean±SEM; NC vs 5xFAD and Vehicle vs SCDi were compared by two-way ANOVA Fisher’s LSD post-hoc test as appropriate (*p<0.05 **p<0.01 ***p<0.001). C16:0, palmitic acid; C16:1, palmitoleic acid; C18:0, stearic acid; C18:1, oleic acid; C18:2, linoleic acid; C20:4, arachidonic acid, C22:6, docosahexaenoic acid; SFA, saturated fatty acids; MUFA, monounsaturated fatty acids; PUFA, polyunsaturated fatty acids.

We conclude that the altered levels of SCD-related fatty acids identified in unoperated 5xFAD female mice remain detectable following pump implantation and vehicle infusion, and that SCDi infusion modulates levels of both SCD-related fatty acids and PUFAs in this more aggressive model of AD pathology.

### Loss of 5xFAD hippocampal dendritic spines is mitigated by SCD inhibition

Lastly, we examined whether the SCDi effects on the 5xFAD hippocampal fatty acid profile are associated with changes in dendritic spine density. Hippocampal synapse loss is a hallmark of AD that is tightly correlated with memory loss and cognitive decline, and previous work showed that SCDi infusion increases the density of postsynaptic dendritic spines in the slow-progressing 3xTg-AD hippocampus. Here, we likewise evaluated its effects on spine density in 5xFAD and NC mice, focusing specifically on the dentate gyrus (DG) and the cornu ammonis 1 (CA1) of the dorsal hippocampus, which is associated with the spatial learning and memory^33^ functions that were improved by SCDi in 3xTg-AD mice^24^.

Neurons were labelled for quantification of secondary and tertiary dendritic spine density using Golgi staining of thick hippocampal sections. In vehicle-infused mice, 5xFAD neurons showed statistically significant decreases of tertiary dendritic spines on both DG granule neurons and CA1 pyramidal neurons (Figure 4), confirming the expected synapse loss in this mouse model of AD. Trends for decreased spine densities on secondary dendrites of these DG and CA1 neurons did not reach statistical significance (Figure 4d and f).

**Figure 4.**
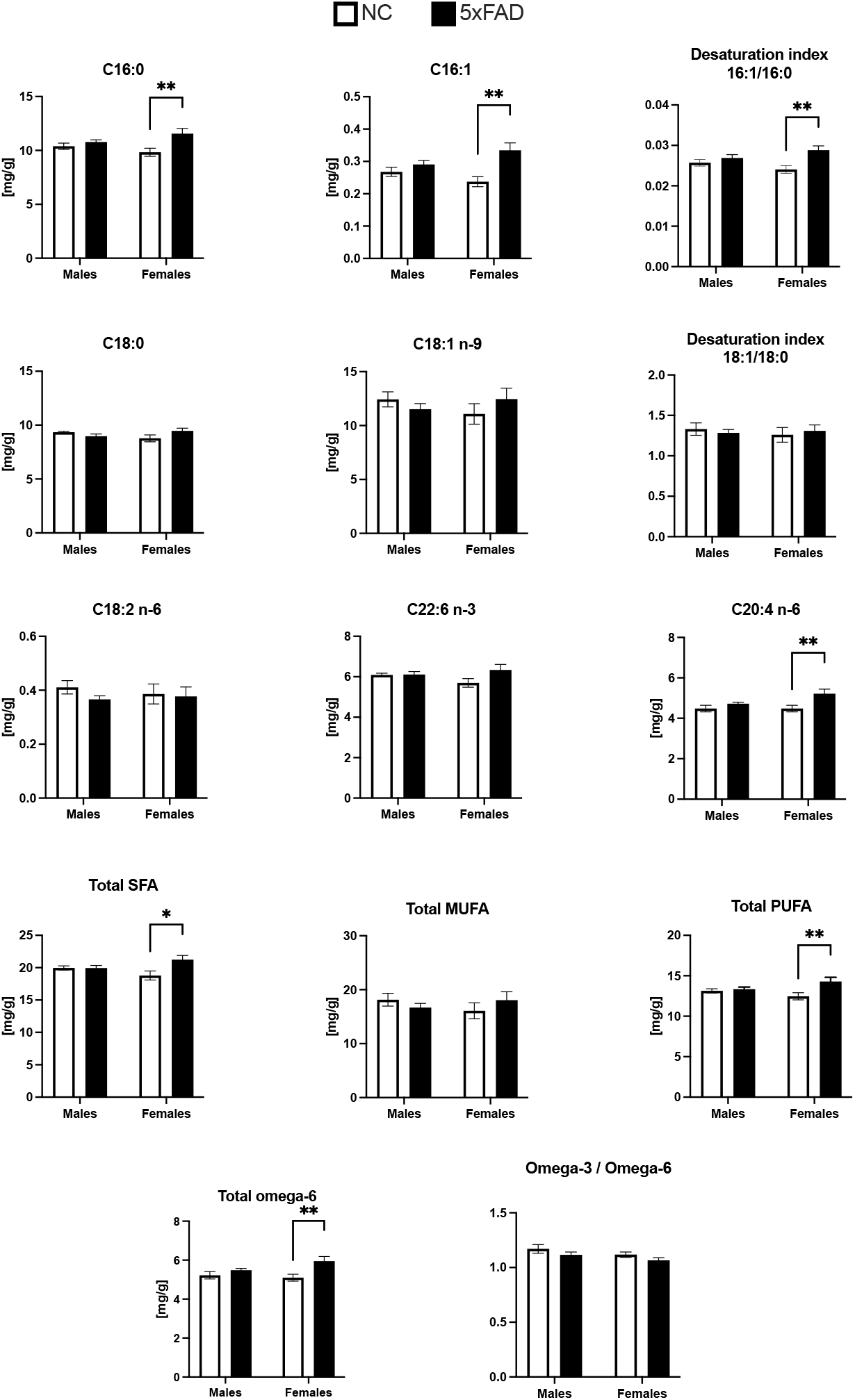
Effects of SCD inhibition on dendritic spines in dentate gyrus (DG) and cornus ammonis 1 (CA1) in 5xFAD. a-c) Golgi staining in the hippocampus. a) Coronal section of the dorsal hippocampus. b-c) Representative pictures of neurons stained with Golgi-Cox staining (4x magnification) and within the regions of interest: b) dentate gyrus (DG) and a granule cell (arrows indicate 2° and 3° dendrites) c) cornu ammonis 1 (CA1) and a pyramidal neuron (arrows indicate 2° and 3° dendrites). d-g) Changes in spine numbers are shown in NC vs 5xFAD, Vehicle vs SCDi-treated female mice of 6 months of age after a month of treatment. d) Dendritic spine density on secondary dendrites of granule cells in the DG and e) on tertiary dendrites in the DG. 2-way ANOVA showed a significant genotype effect with 5xFAD mice having less spine than NC mice (F (1, 105) = 8.445, p=0.0045) but no drug effect or interaction. Dendritic spines were significantly lower in 5xFAD-Vehicle vs NC-Vehicle (p=0.0378) but not following SCDi treatment, NC-Vehicle/5xFAD-SCDi (p=0.3196) f) Dendritic spine density on secondary dendrites of pyramidal neurons in the CA1. 2-way ANOVA showed significant treatment (F (1,105) =18.44, p<0.0001) and interaction effects (F (1,105)=8.414, p=0.0045). Dendritic spines quantification showed no difference in NC-Vehicle/5xFAD-Vehicle (p=0.2627), but a significant increase in spine density following SCDi treatment, NC-Vehicle/5xFAD-SCDi (p=0.0051); and g) on tertiary dendrites. 2-way ANOVA showed a significant genotype effect with 5xFAD mice having less spine than NC mice (F (1, 105) = 7.338, p=0.0079) but no drug effect or interaction. Spines were significantly lower in 5xFAD-Vehicle vs NC-Vehicle (p=0.0099), but not following SCDi treatment, NC-Vehicle/5xFAD-SCDi (p=0.2685). (N=22-37, Two-way ANOVA Dunnett’s post-hoc test *p<0.05 **p<0.01).

In SCDi-infused mice, the loss of tertiary dendritic spines of both DG and CA1 5xFAD neurons was reversed, with these populations exhibiting upward shifts in spine density distributions (Figure 4e and g). Notably, the statistically significant declines in tertiary spine densities of vehicle-infused 5xFAD mice were eliminated in both the DG and CA1 areas of SCDi-infused 5xFAD mice. Furthermore, the secondary spine densities of 5xFAD CA1 neurons were increased to significantly higher levels than found on the vehicle-infused NC mice (Figure 4f).

Collectively, these data reveal age- and sex-related lipidomic changes in the hippocampal fatty acid profile of the 5xFAD model, including in SCD-related fatty acids and PUFA, and reveals that SCD inhibition remains capable of modulating hippocampal fatty acid composition and increasing dendritic spine density in this neuropathologically aggressive AD model.

## Discussion

Our previous work has shown that pharmacological inhibition of a single lipid enzyme, SCD, is sufficient to reverse neurogenic, immune, synaptic and behavioral impairments when administered at pre-pathology timepoints in the slow-progressing 3xTg-AD mouse model^23,24^. Here, we tested whether SCDi can have potential beneficial effects when used in the highly amyloidogenic 5xFAD mouse model, finding that it remains capable of modulating hippocampal fatty acid composition and reversing losses of dendritic spines in the context of rapid and aggressive amyloid plaque formation. Exploiting the 5xFAD model strengthens our prior insights on the effects of AD genetic risk factors on fatty acid metabolism and supports the rationale for developing brain SCD as a pharmacological target.

5xFAD mice overexpress APP (amyloid precursor protein), and PSEN1 (Presenilin 1) containing 5 familial AD mutations (APP KM670/671NL (Swedish), APP I716V (Florida), APP V717I (London), PSEN1 M146L, PSEN1 L286V), under the control of a Thy1 mini-gene, which instructs expression to forebrain neurons^34,35^. These mice develop a rapid and aggressive AD phenotype characterized by robust amyloid plaque pathologies that appear and are well established in the brain between 2–5 months of age. We tested whether the reported microgliosis, inflammatory processes and synaptic loss^28,36,37^ are present within our breeding colony, confirming that amyloid plaques and microgliosis are indeed detectable at 5 months of age. Plaques continued increasing between 5 and 8 months of age in both the cortex and hippocampus, with the hippocampal dentate gyrus appearing more affected than the CA1. Interestingly, Iba1+ microgliosis but not GFAP+ astrogliosis correlated with amyloid plaque pathology, although the lack of obvious morphological changes in GFAP does not exclude the existence of functional changes in astrocytes.

We then extended our characterization of the 5xFAD model to assess fatty acid disturbances. Lipid metabolism alterations are observed in both sporadic and familial AD and support the idea that such disturbances are a common mechanism in all forms of AD^38–40^. Indeed, mechanistically, genome-wide association studies have identified more than 20 AD risk loci, among which there is an enrichment of lipid metabolism and innate immunity genes^5,10,41,42^. This includes the primary genetic risk factor for sporadic AD, APOE4, a gene encoding a lipoprotein involved in triglyceride and cholesterol transport^5^. APOE4 is a risk factor not just for SAD, but also in late-onset FAD^43^, as apoE4 likewise accelerates the development of FAD^44–46^. Characterization of the hippocampus, cortex and plasma fatty acid profiles of 5xFAD mice highlighted alterations within the hippocampus in the absence of changes in the cortex and plasma. Specifically, symptomatic 5xFAD mice displayed increased hippocampal levels of palmitic and palmitoleic acids when compared to the littermate strain controls. Hippocampal SCD-related fatty acid alterations were apparent in 5 MO females and were confirmed in 8 MO females, and also appeared in males at 8 MO, suggesting an aggravation of SCD activity with disease progression. At 5 months of age, the 5xFAD female mice already presented a higher C16:1/C16:0 hippocampal desaturation index versus NC females, an indirect measure of SCD activity^47^, which is reminiscent of what was previously described in the symptomatic females of the 3xTg-AD mouse model^24^. In that previous study, the 3xTg-AD mouse model showed MUFA-rich triglyceride deposits in the subventricular zone that preceded plaques, tangles, or cognitive defects^23^. Here, we extended upon this observation by examining the fatty acid composition of complex lipid classes. This revealed higher C16 desaturation within the hippocampal phospholipid fraction of 5xFAD mice. SCD also uses stearic acid to form oleic acid, but the levels of these FAs remained stable between males and females as well as the C18:1/C18:0 DI, indicating that in this model C18 fatty acids may be buffered or that the palmitic acid SCD-reaction may be selectively affected. Interestingly, little is known about the activity of other fatty acid processing enzymes in AD, such as the elongase ELOVL6, which elongates C16:0 to C18:0 and C16:1 to C18:1 and could therefore be recruited as a coping mechanism for changes in SCD activity. A deeper understanding of the relationship between ELOVL and SCD enzymes could be beneficial to uncover the mechanisms by which FAs are processed.

The cortex is also one of the most affected regions in AD. Total FA concentrations were analyzed in the cortex of 8 MO 5xFAD and NC mice, yet no significant changes were observed. Changes in cortical fatty acid composition have been reported in postmortem human AD brains^21,48,49^ and may require longer to develop in the present model.

Recently, we showed that inhibition of SCD in the 3xTg-AD mouse model leads to rescue of aberrant lipid, immune and synaptic gene expression signatures in the hippocampus, recovery of hippocampal dendritic spines, and rescue of hippocampal-dependent learning and memory^24^ SCD inhibition strategy in the highly amyloidogenic 5xFAD model also modulates the hippocampal fatty acid profile and increases dendritic spine density. Spine density was assessed specifically in the 5xFAD dorsal hippocampus that is dedicated to spatial learning and memory. SCD inhibition rescued the 5xFAD tertiary dendrite spine density of DG and CA1 neurons, which no longer differed between NC-Vehicle and 5xFAD-SCDi mice, and also increased spine density on CA1 secondary dendrites to higher levels than in NC-Vehicle mice.

The precise downstream pathways by which SCD inhibition rescues or regenerates spines *in vivo* merits further investigation. Assessment of the hippocampal fatty acid profile revealed multiple indicators of at least partial inhibition of 5xFAD SCD activity, including significantly increased levels of palmitic acid and total SFA and decreased C18:1/C18:0 desaturation index. Unexpectedly, SCDi also significantly increased total PUFA, including total 22:6 n-3 DHA, 20:4 n-6 AA and total omega-6 fatty acids. Since DHA is the most abundant omega-3 FA in the brain and can be oxidized into anti-inflammatory and neuroprotective metabolites, its increase could be a mediator of the positive effects seen with SCDi treatment^50^. Indeed, DHA is concentrated in synapses and has been implicated in synaptic plasticity, vasculature health, and Aβ production and clearance ^51^. Interestingly, SCDi did not have significant effects on the fatty acid profile of the NC control mice, an observation that is consistent with its potent biological effects on 3xTg-AD mice, but not on their WT controls^24^. Establishing the downstream mechanisms by which SCDi-induced alterations in its target fatty acids leads to reversal of biological impairments in 3xTg-AD and 5xFAD models of AD is a critical next step.

Epidemiological evidence supports sex differences in AD^52^. We observed that male and females 5xFAD mice had similar plaque burden and microgliosis, yet female mice displayed earlier and clearer fatty acid changes than males. We therefore focused our SCDi treatment on 5xFAD females and their NC counterparts, but it will be important to assess whether there are sex-related differences in the response to SCDi, as well as whether the SCDi-mediated lipidomic and dendritic spine effects seen after one month of administration are sufficient to improve cognition in this model as it did in the 3xTg-AD model^24^.

Taken together, these results reinforce the rationale for exploring SCD inhibition as a therapeutic strategy in AD. We demonstrate that the beneficial effects of SCD inhibitor treatment on FA and hippocampal dendritic spines can be extended to a second AD mouse model, and that they remain statistically significant even when administration starts at a stage of heavy brain plaque burden. Recent studies have likewise provided evidence that reducing SCD activity protects against neurodegeneration in models of multiple sclerosis and Parkinson’s disease^25^, encouraging continued clinical development of this promising therapeutic target for neurodegenerative diseases.

## Materials and methods

### Mouse model

5XFAD mice (Jackson laboratory RRID: MMRRC_034840-JAX) were crossed with B6SJLF1/J mice (Jackson laboratory RRID:IMSR_JAX:100012) to have the heterozygous 5xFAD mice and Non-Carrier (NC) non-transgenic littermate controls. All mice were bred in-house, maintained in identical housing conditions (22□°C, 50% humidity, 14h light/ 10h dark cycle) and given free access to a chow feed and water.

Mice were sacrificed at 2, 5 and 8 months of age and brain regions were dissected and stored as appropriate for the analyses. Animal sample size was decided according to previous experiments^24,53^ and to achieve a degree of freedom of ANOVA equal to at least 20. All mice were randomly assigned to experimental groups, ensuring that each litter was distributed across groups.

All experiments were conducted following the guidelines of the Canadian Council of Animal Care (CCAC) and were approved by the Institutional Animal Care Committee (CPA) of the Faculty of Medicine and Health Sciences (FMSS) at the Université de Sherbrooke. All efforts were made to minimize animal suffering and to reduce the number of animals used. Male and female mice were used for this purpose. Mice were sacrificed under deep anesthesia with isoflurane and promptly decapitated for the following experiments. We confirm adherence to the ARRIVE guidelines regarding reporting of results.

### DAB immunohistochemistry

For immunohistochemistry (IHC) 5XFAD and NC 2-5-8 months of age half-brains were post-fixed with paraformaldehyde 4% in phosphate buffer 0.1M (PFA 4%) for 24 hours. The half-brain of each animal was cut into 40 µm coronal sections using a vibrating microtome (Leica VT1000S, Leica Microsystems), and the tissue sections were stored at −20°C in an antifreeze solution. Brightfields IHC was performed as previously detailed^24^ using rabbit anti-Aβ 37-42 (8243S) (Cell Signaling Tech, 1:100), rabbit anti-Iba1 (019-19741) (Wako Chemicals, 1:500) and rabbit anti-GFAP (Z0334) (Dako, 1:500) incubated for an overnight at 4°C. Primary antibodies were detected using Goat anti-rabbit biotinylated secondary antibody (Jackson Immuno Research, 1:1000), and revealed using a 3,3-diaminobenzidine (DAB)-containing solution (Sigma-Aldrich).

### Lipid extraction and analysis of FA profile by gas chromatography–flame ion detection (GC–FID)

Plasma, hippocampus and cortex of 5XFAD and NC 2, 5 and 8 MO were collected to analyze total FA profile and FA profile of TG, PL, CE and FFA. Total lipids were extracted from plasma, hippocampus and cortex samples using the Folch method^54^ and prior to extraction internal standard (iSTD) for total fatty acid, C17:0 PC, was added in amounts depending on the sample (0.05 mg iSTD per 0.01 g hippocampus, 0.1 mg per 0.01 mg cortex); and the iSTD C17:0 TG 0.12 mg per 50µL of plasma. iSTD concentration calculations for each lipid class were based on lipids concentrations in plasma according to Quehnberger et al. and in hippocampus according to Fitzner et al.^55,56^ (Supplementary Table S7). Tissues were then homogenized using the Bead Mill Homogenizer (VWR, cat# 75840-022). The lipid extract was evaporated under nitrogen and reconstituted with 200 μL of chloroform.

To separate TG, PL, CE and FFA, the lipid extract was collected and loaded in Bond Elut NH2 200mg cartridges (Agilent Technologies). Chloroforme:isopropanol (2:1 v/v) solution was added twice to each cartridge for elution of neutral lipids (TG and CE)^57^. Ethyl eter:acetic acid (98.7:1.3 v/v) solution was added for subsequent elution of FFA. Methanol was added for elution of PL. Neutral lipid fraction underwent to a second cartridge extraction for the separation of TG and CE after solvents evaporation. This fraction was loaded and eluted twice with hexane to collect the CE and twice with hexane:diethyl ether:MeCl2 (1%-10% v/v) to collect the TG^58^

Each extracted sample underwent to saponification to release the fatty acid salts from each lipid molecule. Saponification of the fatty acids was performed using KOH-methanol and followed by protonation with 12 N HCl and methylation with 12% BF3-methanol (Thermo Fisher Scientific cat# AC402765000), as described^59^.

The total fatty acid composition and fatty acid composition of each lipid class was analyzed by gas chromatography coupled with a flame ionization detector (model 6890; Agilent). 1 μl of the sample was injected in splitless mode at 250 °C. The fatty acid methyl esters (FAME) were carried by helium at a pressure of 107 kPA through a 50m BPX-70 fused capillary column (0.22-mm diameter, 0.25-μm film thickness; SGE). A thermal gradient was applied on the column. FAMEs were then detected at the end of the column by the flame ionization detector at 260 °C. The chromatogram analysis was performed using OpenLab CDS ChemStation version C01.10.

### In vivo surgical procedures

5xFAD and NC female mice 5 MO were locally injected with bupivacaine and operated under isoflurane anesthesia. Intracerebroventricular Alzet 28-day osmotic pumps (0.11µl/h infusion rate, model 1004; Durect, cat# 0009922) linked to brain infusion cannula (Durect, cat# 0008852) were stereotaxically implanted into the left lateral ventricle (coordinates: 0mm AP and -0.9mm ML to the bregma) and the pumps placed under the back skin according to manufacturer’s instructions. The pumps contained the vehicle, DMSO (Sigma-Aldrich, cat# D2650), or 80 µM SCD1 inhibitor (Abcam, cat# ab142089) diluted in DMSO. Both were previously diluted in artificial cerebrospinal fluid (148□mM NaCl, 3□mM KCl, 1.7□mM MgCl2, 1.4□mM CaCl2, 1.5□mM Na2PO4, 0.1□mM NaH2PO4) following the manufacturer’s instructions.

### Dendritic spines quantification (Golgi staining)

For hippocampal dendritic spines quantification, one brain hemisphere was cut in a mouse brain mold at 4mm from the cerebellum and 5mm from the olfactory bulb. The staining was performed with the slice Golgi kit (Bioenno Tech, cat# 003760) following the manufacturer’s instructions. Briefly, half-brains were post-fixed and sliced on a vibratome to obtain 12 sections of 150µm and incubated in 0.1M PB pH 7.4, washed with the same solution and incubated in impregnation solution for 6 days at room temperature. The sections were washed in water and 3 times in phosphate buffer saline – Triton X (PBS-T) pH 7.4 and 2 times in 0.01M PBS pH 7.4. The Golgi staining was performed by adding solution C from the kit, followed by incubation in solution D. Subsequent washes were performed in PBS before dehydrating the sections in increasing concentrations of ethanol of 50%, 80%, 95%, and 100%, followed by two washes in xylene. The sections were mounted on slides with Permount (Thermofisher Scientific) and left to dry under a chemical hood for 48 hours before imaging and quantification.

For the quantification of spines, two neurons in the dentate gyrus (DG) and two neurons in the cornu ammonis 1 (CA1) of the dorsal hippocampi were selected for quantification. The neurons had to have a visible cell body as well as secondary and tertiary dendrites to be selected for quantification. The dorsal hippocampus sections were between -1.06mm to - 1.94mm from Bregma. The dendritic spine density was determined by counting the spines on the secondary and tertiary dendrites of the neurons on a length of 19µm, using a brightfield microscope at 100X magnification (Olympus BX43). Dendritic spines were quantified on coded slides by a blinded observer.

### Statistical analyses

Statistical analyses were done using GraphPad Prism 10.0 (GraphPad Software). Data are presented as means ± standard errors of the mean. Each statistical test performed is detailed in the legend of the figures and P-values smaller than 0.05 were considered statistically significant.

## Data Availability

The data generated during this study are available from the corresponding author upon reasonable request.

## Author contributions statement

MT co-wrote the manuscript. MT, MA carried out experiments in collaboration with AA, AV, JAL, MM. MT analyzed data and prepared figures. AV and MP provided training and quality control for GC-FID analysis. KF analyzed data, co-wrote the manuscript, and developed the concept. All authors reviewed the manuscript.

## Additional information

The authors declare that there are no competing interests.

## References

1. A Armstrong, R. Risk factors for Alzheimer’s disease. Folia Neuropathol 57, 87–105 (2019).

2. Launer, L. J., White, L. R., Petrovitch, H., Ross, G. W. & Curb, J. D. Cholesterol and neuropathologic markers of AD: a population-based autopsy study. Neurology 57, 1447–52 (2001).

3. Shi, Y. & Holtzman, D. M. Interplay between innate immunity and Alzheimer disease: APOE and TREM2 in the spotlight. Nat Rev Immunol 18, 759–772 (2018).

4. Nguyen, A. T. et al. APOE and TREM2 regulate amyloid-responsive microglia in Alzheimer’s disease. Acta Neuropathol 140, 477–493 (2020).

5. Kunkle, B. W. et al. Genetic meta-analysis of diagnosed Alzheimer’s disease identifies new risk loci and implicates Aβ, tau, immunity and lipid processing. Nat Genet 51, 414–430 (2019).

6. Turri, M., Marchi, C., Adorni, M. P., Calabresi, L. & Zimetti, F. Emerging role of HDL in brain cholesterol metabolism and neurodegenerative disorders. Biochim Biophys Acta Mol Cell Biol Lipids 1867, 159123 (2022).

7. Farrer, L. A. et al. Effects of age, sex, and ethnicity on the association between apolipoprotein E genotype and Alzheimer disease. A meta-analysis. APOE and Alzheimer Disease Meta Analysis Consortium. JAMA 278, 1349–56.

8. Neu, S. C. et al. Apolipoprotein E Genotype and Sex Risk Factors for Alzheimer Disease: A Meta-analysis. JAMA Neurol 74, 1178–1189 (2017).

9. Jansen, I. E. et al. Genome-wide meta-analysis identifies new loci and functional pathways influencing Alzheimer’s disease risk. Nat Genet 51, 404– 413 (2019).

10. Lambert, J. C. et al. Meta-analysis of 74,046 individuals identifies 11 new susceptibility loci for Alzheimer’s disease. Nat Genet 45, 1452–8 (2013).

11. Karch, C. M. & Goate, A. M. Alzheimer’s disease risk genes and mechanisms of disease pathogenesis. Biol Psychiatry 77, 43–51 (2015).

12. Heath, L. et al. Manifestations of Alzheimer’s disease genetic risk in the blood are evident in a multiomic analysis in healthy adults aged 18 to 90. Sci Rep 12, 6117 (2022).

13. Kivipelto, M. et al. World-Wide FINGERS Network: A global approach to risk reduction and prevention of dementia. Alzheimers Dement 16, 1078–1094 (2020).

14. Rosenberg, A., Mangialasche, F., Ngandu, T., Solomon, A. & Kivipelto, M. Multidomain Interventions to Prevent Cognitive Impairment, Alzheimer’s Disease, and Dementia: From FINGER to World-Wide FINGERS. J Prev Alzheimers Dis 7, 29–36 (2020).

15. Livingston, G. et al. Dementia prevention, intervention, and care: 2020 report of the Lancet Commission. Lancet 396, 413–446 (2020).

16. Scheltens, P. et al. Alzheimer’s disease. Lancet 397, 1577–1590 (2021).

17. Edwards Iii, G. A., Gamez, N., Escobedo, G., Calderon, O. & Moreno-Gonzalez, I. Modifiable Risk Factors for Alzheimer’s Disease. Front Aging Neurosci 11, 146 (2019).

18. Huynh, K. et al. Concordant peripheral lipidome signatures in two large clinical studies of Alzheimer’s disease. Nat Commun 11, (2020).

19. Snigdha, S., Astarita, G., Piomelli, D. & Cotman, C. W. Effects of diet and behavioral enrichment on free fatty acids in the aged canine brain. Neuroscience 202, 326–33 (2012).

20. Fraser, T., Tayler, H. & Love, S. Fatty acid composition of frontal, temporal and parietal neocortex in the normal human brain and in Alzheimer’s disease. Neurochem Res 35, 503–13 (2010).

21. Cunnane, S. C. et al. Plasma and brain fatty acid profiles in mild cognitive impairment and Alzheimer’s disease. J Alzheimers Dis 29, 691–7 (2012).

22. Astarita, G. et al. Elevated stearoyl-CoA desaturase in brains of patients with Alzheimer’s disease. PLoS One 6, e24777 (2011).

23. Hamilton, L. K. et al. Aberrant Lipid Metabolism in the Forebrain Niche Suppresses Adult Neural Stem Cell Proliferation in an Animal Model of Alzheimer’s Disease. Cell Stem Cell 17, 397–411 (2015).

24. Hamilton, L. K. et al. Stearoyl-CoA Desaturase inhibition reverses immune, synaptic and cognitive impairments in an Alzheimer’s disease mouse model. Nat Commun 13, (2022).

25. Loix, M. et al. Stearoyl-CoA desaturase-1: a potential therapeutic target for neurological disorders. Mol Neurodegener 19, 85 (2024).

26. Bogie, J. F. J. et al. Stearoyl-CoA desaturase-1 impairs the reparative properties of macrophages and microglia in the brain. J Exp Med 217, (2020).

27. Fanning, S. et al. Lipidomic Analysis of α-Synuclein Neurotoxicity Identifies Stearoyl CoA Desaturase as a Target for Parkinson Treatment. Mol Cell 73, 1001-1014.e8 (2019).

28. Forner, S. et al. Systematic phenotyping and characterization of the 5xFAD mouse model of Alzheimer’s disease. Sci Data 8, 270 (2021).

29. Oblak, A. L. et al. Comprehensive Evaluation of the 5XFAD Mouse Model for Preclinical Testing Applications: A MODEL-AD Study. Front Aging Neurosci 13, 713726 (2021).

30. Klawitter, J. et al. Fatty acid desaturation index in human plasma: comparison of different analytical methodologies for the evaluation of diet effects. Anal Bioanal Chem 406, 6399–408 (2014).

31. Yoon, J. H. et al. Brain lipidomics: From functional landscape to clinical significance. Sci Adv 8, (2022).

32. Kumar, M. et al. Triglycerides are an important fuel reserve for synapse function in the brain. Nat Metab 1–12 (2025) doi:10.1038/S42255-025-01321-X;SUBJMETA=287,319,378,443,45,631,87;KWRD=CELLULAR+NEUROSCIENCE,LIPIDS,METABOLISM.

33. Fanselow, M. S. & Dong, H.-W. Are the dorsal and ventral hippocampus functionally distinct structures? Neuron 65, 7–19 (2010).

34. Moechars, D., Lorent, K., De Strooper, B., Dewachter, I. & Van Leuven, F. Expression in brain of amyloid precursor protein mutated in the alpha-secretase site causes disturbed behavior, neuronal degeneration and premature death in transgenic mice. EMBO J 15, 1265–74 (1996).

35. Vidal, M., Morris, R., Grosveld, F. & Spanopoulou, E. Tissue-specific control elements of the Thy-1 gene. EMBO J 9, 833–40 (1990).

36. Oakley, H. et al. Intraneuronal beta-amyloid aggregates, neurodegeneration, and neuron loss in transgenic mice with five familial Alzheimer’s disease mutations: potential factors in amyloid plaque formation. J Neurosci 26, 10129–40 (2006).

37. Jawhar, S., Trawicka, A., Jenneckens, C., Bayer, T. A. & Wirths, O. Motor deficits, neuron loss, and reduced anxiety coinciding with axonal degeneration and intraneuronal Aβ aggregation in the 5XFAD mouse model of Alzheimer’s disease. Neurobiol Aging 33, 196.e29–40 (2012).

38. Kao, Y. C., Ho, P. C., Tu, Y. K., Jou, I. M. & Tsai, K. J. Lipids and alzheimer’s disease. Int J Mol Sci 21, (2020).

39. Wong, M. W. et al. Dysregulation of lipids in Alzheimer’s disease and their role as potential biomarkers. Alzheimer’s and Dementia 13, 810–827 (2017).

40. Turri, M. et al. Plasma and cerebrospinal fluid cholesterol esterification is hampered in Alzheimer’s disease. Alzheimers Res Ther 15, (2023).

41. Kunkle, B. W. et al. Novel Alzheimer Disease Risk Loci and Pathways in African American Individuals Using the African Genome Resources Panel: A Meta-analysis. JAMA Neurol 78, 102–113 (2021).

42. Olgiati, P., Politis, A. M., Papadimitriou, G. N., De Ronchi, D. & Serretti, A. Genetics of late-onset Alzheimer’s disease: update from the alzgene database and analysis of shared pathways. Int J Alzheimers Dis 2011, 832379 (2011).

43. Dorszewska, J., Prendecki, M., Oczkowska, A., Dezor, M. & Kozubski, W. Molecular Basis of Familial and Sporadic Alzheimer’s Disease. Curr Alzheimer Res 13, 952–63 (2016).

44. Wijsman, E. M. et al. APOE and other loci affect age-at-onset in Alzheimer’s disease families with PS2 mutation. Am J Med Genet B Neuropsychiatr Genet 132B, 14–20 (2005).

45. Ryman, D. C. et al. Symptom onset in autosomal dominant Alzheimer disease: a systematic review and meta-analysis. Neurology 83, 253–60 (2014).

46. Pastor, P. et al. Apolipoprotein Eepsilon4 modifies Alzheimer’s disease onset in an E280A PS1 kindred. Ann Neurol 54, 163–9 (2003).

47. Klawitter, J. et al. Fatty acid desaturation index in human plasma: comparison of different analytical methodologies for the evaluation of diet effects. Anal Bioanal Chem 406, 6399–408 (2014).

48. Prasad, M. R., Lovell, M. A., Yatin, M., Dhillon, H. & Markesbery, W. R. Regional membrane phospholipid alterations in Alzheimer’s disease. Neurochem Res 23, 81–8 (1998).

49. Martín, V. et al. Lipid alterations in lipid rafts from Alzheimer’s disease human brain cortex. J Alzheimers Dis 19, 489–502 (2010).

50. McNamara, R. K. & Carlson, S. E. Role of omega-3 fatty acids in brain development and function: potential implications for the pathogenesis and prevention of psychopathology. Prostaglandins Leukot Essent Fatty Acids 75, 329–49 (2006).

51. Yassine, H. N. et al. Baseline Findings of PreventE4: A Double-Blind Placebo Controlled Clinical Trial Testing High Dose DHA in APOE4 Carriers before the Onset of Dementia. J Prev Alzheimers Dis 10, 810–820 (2023).

52. Aggarwal, N. T. & Mielke, M. M. Sex Differences in Alzheimer’s Disease. Neurol Clin 41, 343–358 (2023).

53. Hamilton, L. K. et al. Central inhibition of stearoyl-CoA desaturase has minimal effects on the peripheral metabolic symptoms of the 3xTg Alzheimer’s disease mouse model. Sci Rep 14, (2024).

54. Folch, J., Lees, M. & Sloane Stanley, G. H. A SIMPLE METHOD FOR THE ISOLATION AND PURIFICATION OF TOTAL LIPIDES FROM ANIMAL TISSUES. Journal of Biological Chemistry 226, 497–509 (1957).

55. Quehenberger, O. et al. Lipidomics reveals a remarkable diversity of lipids in human plasma1. J Lipid Res 51, 3299–3305 (2010).

56. Fitzner, D. et al. Cell-Type- and Brain-Region-Resolved Mouse Brain Lipidome. Cell Rep 32, (2020).

57. Chouinard-Watkins, R. et al. Interaction between BMI and APOE genotype is associated with changes in the plasma long-chain-PUFA response to a fish-oil supplement in healthy participants. American Journal of Clinical Nutrition 102, 505–513 (2015).

58. Kaluzny, M. A., Duncan, L. A., Merritt, M. V. & Epps, D. E. Rapid separation of lipid classes in high yield and purity using bonded phase columns. J Lipid Res 26, 135–140 (1985).

59. Chevalier, L., Vachon, A. & Plourde, M. D. S. Pharmacokinetics of Supplemental Omega-3 Fatty Acids Esterified in Monoglycerides, Ethyl Esters, or Triglycerides in Adults in a Randomized Crossover Trial. Journal of Nutrition 151, 1111–1118 (2021).

