## Supplemental for "In vivo inhibition of stearoyl-CoA desaturase modulates the hippocampal fatty acid profile and restores density of dendritic spines in the aggressive 5xFAD model of Alzheimer’s disease"

#### Title:

**Correspondance to:**

*Karl Fernandes, PhD

### Supplementary data

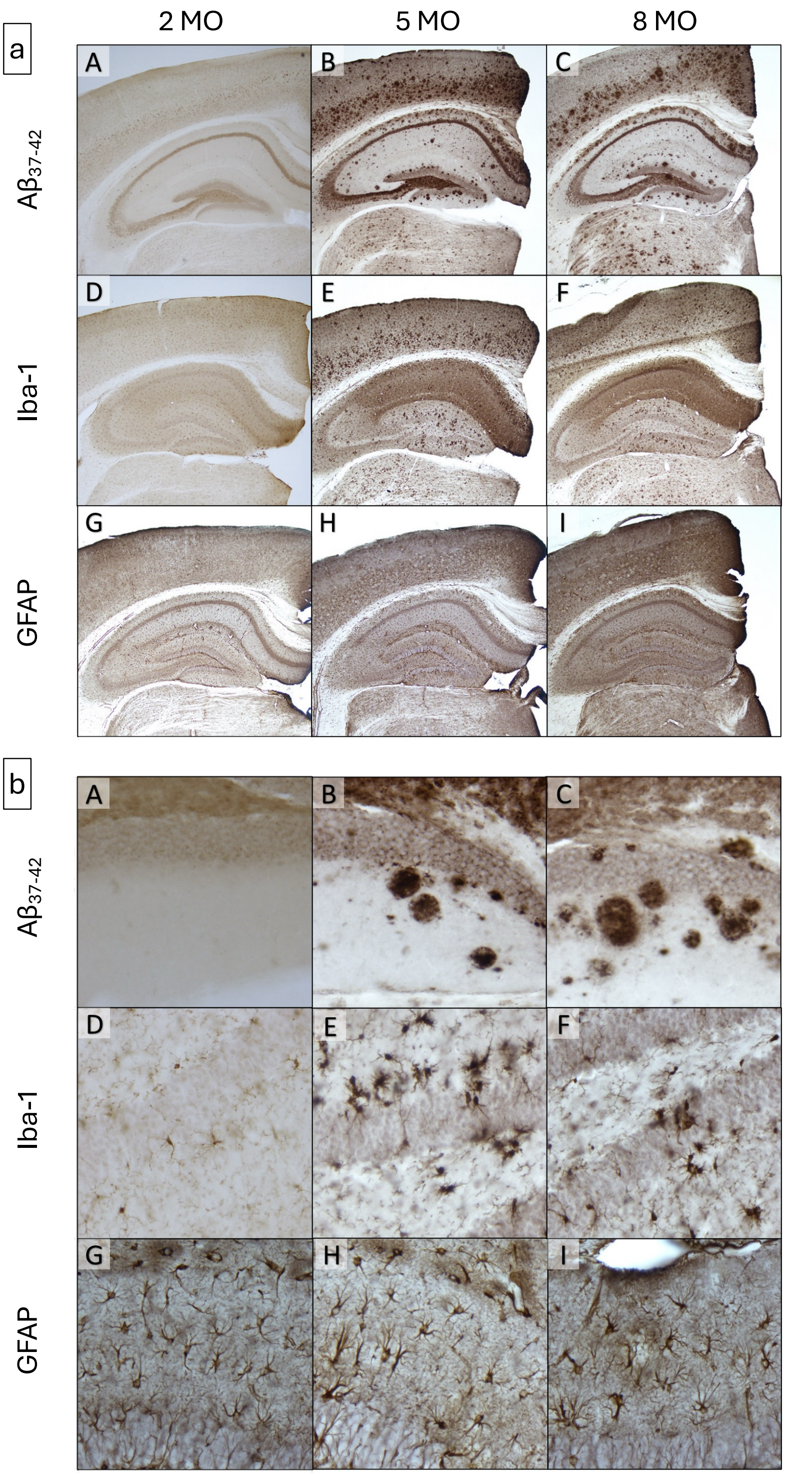


**Supplementary figure S1. Cerebral amyloid deposition and gliosis in the hippocampus of 5xFAD.** Representative 4X (a) and 40X (b) images of 5xFAD male and female mice: (A-C) Aβ37-

42 immunohistochemistry of amyloid plaques in the hippocampus and cortex at 2 months old (A), 5 months old (B), and 8 months old (C). (D-F) Immunohistochemistry of microglia with Iba-1 marker in the hippocampus and cortex at 2 months old (D), 5 months old (E), and 8 months old (F). (G-I) Immunohistochemistry of astrocytes with GFAP marker in the hippocampus and cortex at 2 months old (G), 5 months old (H), and 8 months old (I). (N=4)

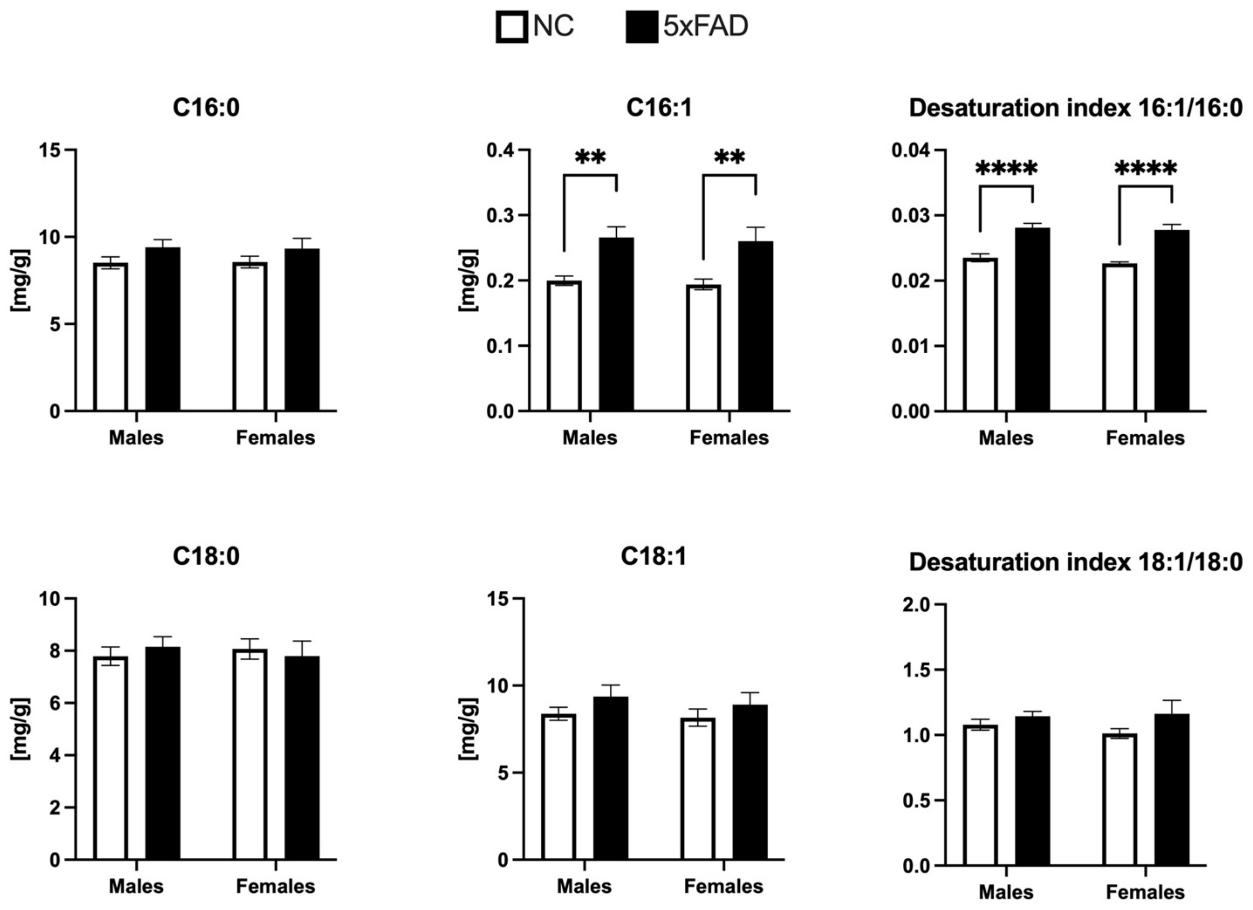


**Supplementary figure S2. Hippocampus total fatty acid composition of 5xFAD at 8 months of age.** Column graph represents the specific fatty acid amount in hippocampus of 5xFAD (black columns, N=4) and NC (white columns, N=4) at 8 months of age. The most relevant hippocampus fatty acids are shown in male and female mice. Desaturation indexes (DI) of palmitoleic over palmitic acid, and oleic over stearic are displayed. Omega-6/omega-3 ratio is shown. Data are expressed as mean±SEM; 5xFAD vs NC were compared by two-way ANOVA with with Šídák's multiple comparisons test as appropriate (**p<0.01, ****p<0.0001). C16:0, palmitic acid; C16:1, palmitoleic acid; C18:0, stearic acid; C18:1, oleic acid.

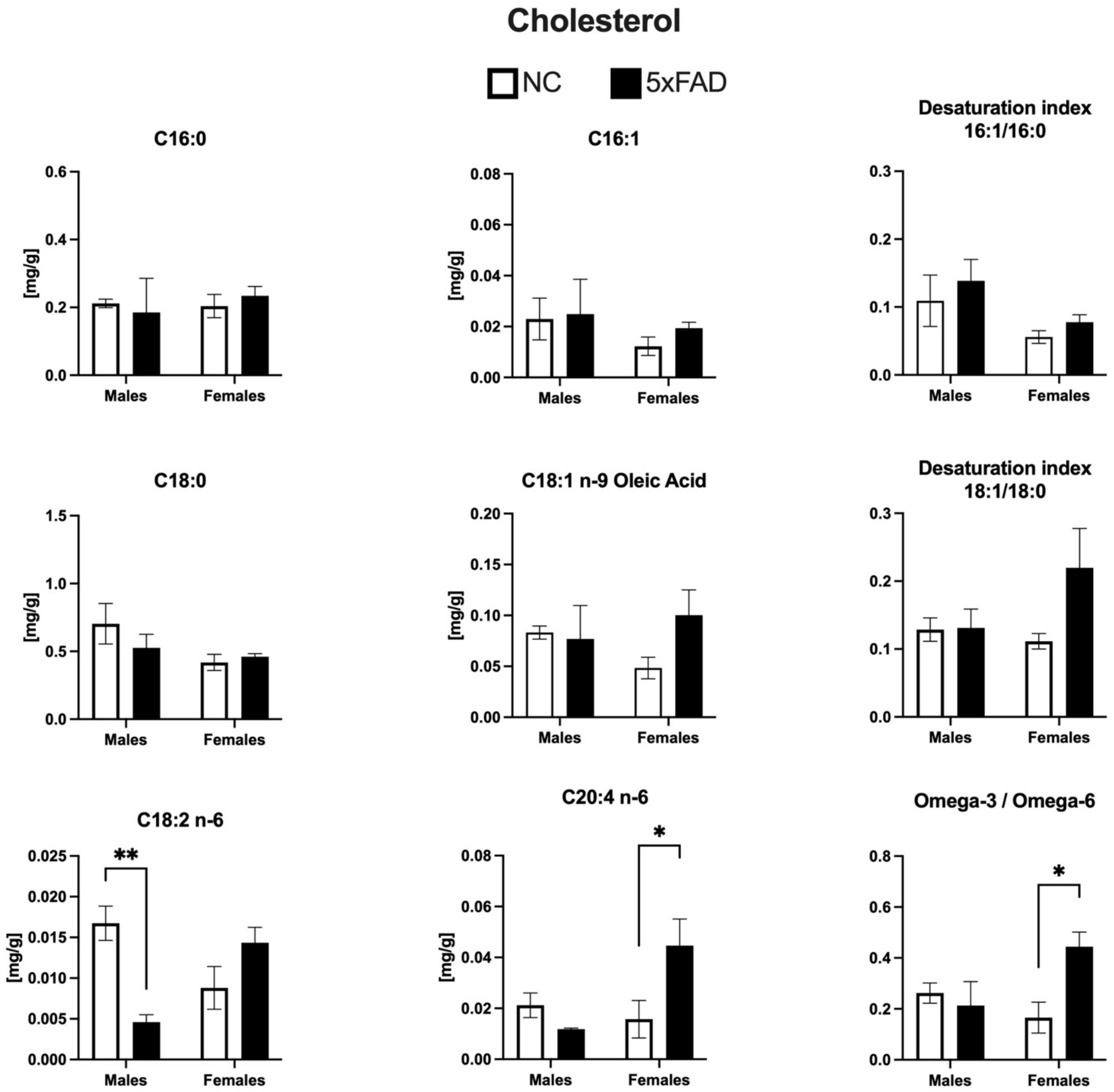


**Supplementary figure S3. Hippocampus cholesteryl esters fatty acid composition of 5xFAD.** Column graph represents the specific cholesteryl esters fatty acid amount in hippocampus of 5xFAD (black columns, N=4) and NC (white columns, N=4) at 5 months of age. The most relevant hippocampus fatty acids are shown in male and female mice. Desaturation indexes (DI) of palmitoleic over palmitic acid, and oleic over stearic are displayed. Omega-6/omega-3 ratio is shown. Data are expressed as mean±SEM; 5xFAD vs NC were compared by two-way ANOVA with with Šídák's multiple comparisons test as appropriate (*p<0.05 **p<0.01). C16:0, palmitic acid; C16:1, palmitoleic acid; C18:0, stearic acid; C18:1, oleic acid; C18:2, linoleic acid; C20:4, arachidonic acid.

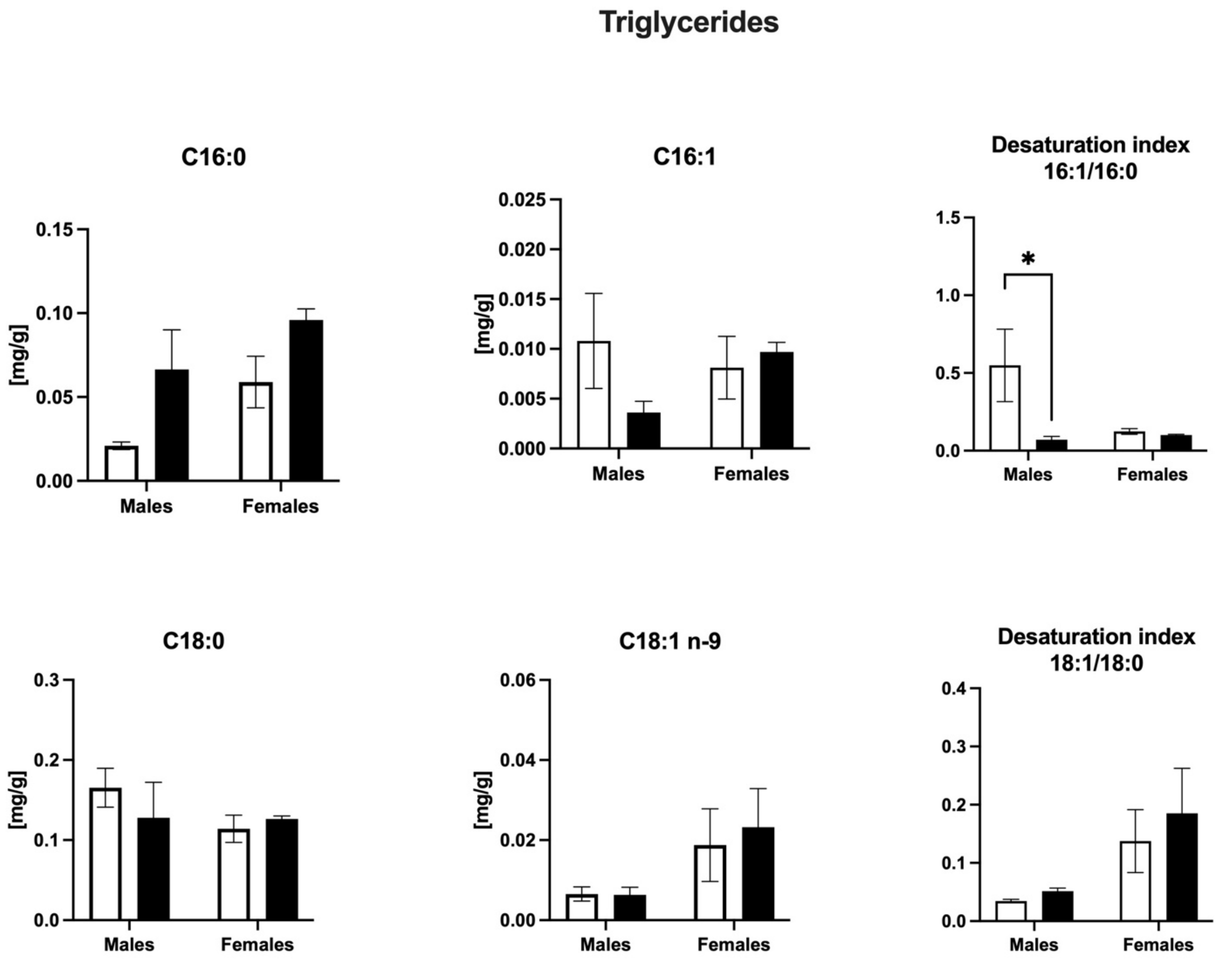


**Supplementary figure S4. Hippocampus triglycerides fatty acid composition of 5xFAD.** Column graph represents the specific fatty acid amount in hippocampus of 5xFAD (black columns, N=4) and NC (white columns, N=4) at 5 months of age. The most relevant hippocampus triglycerides fatty acids are shown in male and female mice. Desaturation indexes (DI) of palmitoleic over palmitic acid, and oleic over stearic are displayed. Data are expressed as mean±SEM; 5xFAD vs NC were compared by two-way ANOVA with with Šídák's multiple comparisons test as appropriate (*p<0.05 **p<0.01). C16:0, palmitic acid; C16:1, palmitoleic acid; C18:0, stearic acid; C18:1, oleic acid.

#### Supplementary Tables

| Fatty acid | 5 MO | | | | | 8 MO | | | | |
|---|---|---|---|---|---|---|---|---|---|---|
| | NC | | 5xFAD | | p value | NC | | 5xFAD | | p value |
| | Mean | SD | Mean | SD | | Mean | SD | Mean | SD | |
| C14:0 | 0,09 | 0,01 | 0,09 | 0,01 | >0.9999 | 0,04 | 0,02 | 0,07 | 0,01 | 0.0409 |
| C16:0 | 10,39 | 0,58 | 10,79 | 0,47 | 0.7314 | 8,51 | 0,85 | 9,41 | 1,05 | 0.2980 |
| C16:1 t | 0,07 | 0,01 | 0,09 | 0,01 | 0.1622 | 0,05 | 0,01 | 0,08 | 0,02 | 0.0402 |
| C16:1 | 0,27 | 0,03 | 0,29 | 0,03 | 0.629 | 0,20 | 0,02 | 0,27 | 0,04 | 0.0083 |
| C18:0 | 9,34 | 0,17 | 8,97 | 0,53 | 0.5678 | 7,79 | 0,87 | 8,15 | 0,94 | 0.8101 |
| C18:1 n-9 | 12,43 | 1,40 | 11,52 | 1,30 | 0.7397 | 8,38 | 0,92 | 9,37 | 1,61 | 0.4069 |
| C18:1 n-7 | 1,58 | 0,09 | 1,58 | 0,12 | >0.9999 | 1,29 | 0,14 | 1,47 | 0,22 | 0.2231 |
| C18:2 t | 0,85 | 0,09 | 0,75 | 0,13 | 0.5870 | 0,51 | 0,10 | 0,50 | 0,18 | 0.9887 |
| C18:2 n-6 | 0,41 | 0,05 | 0,37 | 0,03 | 0.5674 | 0,25 | 0,05 | 0,40 | 0,17 | 0.0601 |
| C20:0 | 0,12 | 0,01 | 0,10 | 0,01 | 0.1688 | 0,05 | 0,04 | 0,09 | 0,02 | 0.1845 |
| C20:1 | 0,97 | 0,10 | 0,83 | 0,15 | 0.5571 | 0,62 | 0,11 | 0,66 | 0,13 | 0.8196 |
| C20:3 n-6 | 0,18 | 0,01 | 0,21 | 0,03 | 0.4473 | 0,16 | 0,02 | 0,24 | 0,09 | 0.1376 |
| C20:4 n-6 | 4,48 | 0,33 | 4,72 | 0,18 | 0.5784 | 4,14 | 0,45 | 4,52 | 0,45 | 0.3327 |
| C22:3 | 0,96 | 0,03 | 1,01 | 0,06 | 0.6707 | 0,94 | 0,11 | 0,96 | 0,12 | 0.9556 |
| C22:5 n-3 | 0,15 | 0,02 | 0,18 | 0,03 | 0.3578 | 0,12 | 0,05 | 0,11 | 0,06 | 0.8906 |
| C22:6 n-3 | 6,09 | 0,17 | 6,11 | 0,35 | 0.9978 | 5,69 | 0,66 | 5,41 | 0,39 | 0.6710 |
| FA tot | 51,27 | 3,11 | 50,05 | 3,37 | 0.9091 | 39,83 | 3,73 | 43,21 | 5,02 | 0.4392 |

**Supplementary Table S1. Hippocampal total fatty acids in 5xFAD male mice at 5 and 8 months of age.** Data are expressed as mean±SEM; 5xFAD vs NC were compared by two-way ANOVA with with Šídák's multiple comparisons test as appropriate.

| Fatty acid | 5 MO | | | | | 8 MO | | | | |
|---|---|---|---|---|---|---|---|---|---|---|
| | NC | | 5xFAD | | P value | NC | | 5xFAD | | P value |
| | Mean | SD | Mean | SD | | Mean | SD | Mean | SD | |
| C14:0 | 0,07 | 0,01 | 0,10 | 0,03 | 0.0159 | 0,05 | 0,01 | 0,07 | 0,02 | 0.1025 |
| C16:0 | 9,83 | 0,92 | 11,56 | 1,17 | 0.0051 | 8,56 | 0,83 | 9,33 | 1,45 | 0.4038 |
| C16:1 t | 0,07 | 0,01 | 0,10 | 0,01 | <0.0001 | 0,05 | 0,02 | 0,08 | 0,02 | 0.0618 |
| C16:1 | 0,24 | 0,04 | 0,33 | 0,06 | 0.0012 | 0,19 | 0,02 | 0,26 | 0,05 | 0.0080 |
| C18:0 | 8,78 | 0,78 | 9,47 | 0,61 | 0.1142 | 8,07 | 0,95 | 7,79 | 1,41 | 0.8833 |
| C18:1 n-9 | 11,08 | 2,32 | 12,46 | 2,46 | 0.4316 | 8,16 | 1,21 | 8,91 | 1,67 | 0.5925 |
| C18:1 n-7 | 1,45 | 0,14 | 1,75 | 0,20 | 0.0056 | 1,32 | 0,14 | 1,41 | 0,23 | 0.6407 |
| C18:2 t | 0,72 | 0,17 | 0,87 | 0,23 | 0.2784 | 0,50 | 0,10 | 0,52 | 0,13 | 0.9311 |
| C18:2 n-6 | 0,39 | 0,09 | 0,38 | 0,09 | 0.9703 | 0,29 | 0,10 | 0,29 | 0,07 | 0.9993 |
| C20:0 | 0,09 | 0,05 | 0,11 | 0,04 | 0.9709 | 0,06 | 0,04 | 0,06 | 0,05 | 0.9937 |
| C20:1 | 0,82 | 0,21 | 0,93 | 0,29 | 0.6155 | 0,60 | 0,12 | 0,59 | 0,13 | 0.9934 |
| C20:3 n-6 | 0,13 | 0,06 | 0,21 | 0,02 | 0.0033 | 0,10 | 0,09 | 0,15 | 0,07 | 0.5687 |
| C20:4 n-6 | 4,48 | 0,40 | 5,22 | 0,55 | 0.0093 | 4,31 | 0,45 | 4,44 | 0,58 | 0.8847 |
| C22:3 | 0,95 | 0,12 | 1,14 | 0,06 | 0.0010 | 0,97 | 0,13 | 0,92 | 0,13 | 0.7354 |
| C22:5 n-3 | 0,10 | 0,05 | 0,14 | 0,01 | 0.1206 | 0,05 | 0,05 | 0,04 | 0,06 | 0.9835 |
| C22:6 n-3 | 5,70 | 0,52 | 6,34 | 0,67 | 0.0704 | 5,70 | 0,59 | 5,21 | 0,71 | 0.3188 |
| FA tot | 47,35 | 5,57 | 53,76 | 5,96 | 0.0653 | 39,96 | 4,45 | 41,30 | 6,26 | 0.8736 |

**Supplementary Table S2. Hippocampal total fatty acids in 5xFAD female mice at 5 and 8 months of age.** Data are expressed as mean±SEM; 5xFAD vs NC were compared by two-way ANOVA with with Šídák's multiple comparisons test as appropriate.

| | 2 MO | | | | | 5 MO | | | | | 8 MO | | | | |
|---|---|---|---|---|---|---|---|---|---|---|---|---|---|---|---|
| | NC | | 5xFAD | | p value | NC | | 5xFAD | | p value | NC | | 5xFAD | | p value |
| **Fatty acid** | Mean | SD | Mean | SD | | Mean | SD | Mean | SD | | Mean | SD | Mean | SD | |
| C14:0 | 0,39 | 0,19 | 4,91 | 8,84 | >0.9999 | 0,40 | 0,20 | 0,58 | 0,28 | 0.5653 | 0,83 | 0,40 | 0,59 | 0,12 | 0.7650 |
| C16:0 | 63,99 | 7,15 | 87,26 | 44,79 | 0.9943 | 60,04 | 16,14 | 64,80 | 9,82 | 0.9406 | 73,20 | 23,77 | 64,86 | 14,16 | 0.4149 |
| C16:1 t | 1,34 | 0,24 | 1,12 | 0,76 | >0.9999 | 1,69 | 0,63 | 1,74 | 0,59 | 0.9994 | 2,11 | 0,53 | 1,83 | 0,66 | 0.8884 |
| C16:1 | 4,97 | 1,03 | 6,51 | 1,99 | 0.9986 | 5,70 | 2,70 | 6,09 | 1,51 | 0.9908 | 5,94 | 3,49 | 4,92 | 1,82 | 0.9170 |
| C18:0 | 39,35 | 2,15 | 47,11 | 14,00 | 0.9982 | 36,72 | 5,71 | 36,37 | 3,54 | 0.9991 | 43,83 | 7,52 | 46,52 | 8,27 | 0.9995 |
| C18:1 n-9 | 38,61 | 3,57 | 52,88 | 27,63 | 0.9858 | 46,28 | 22,65 | 47,83 | 6,80 | 0.9992 | 49,35 | 16,88 | 43,53 | 17,00 | 0.9292 |
| C18:1 n-7 | 6,26 | 0,60 | 7,43 | 2,97 | 0.9988 | 8,97 | 2,72 | 8,78 | 1,39 | 0.9995 | 8,23 | 2,79 | 8,44 | 2,77 | 0.9769 |
| C18:2 n-6 | 90,98 | 13,87 | 91,86 | 21,28 | >0.9999 | 95,85 | 19,00 | 94,97 | 13,36 | >0.9999 | 109,43 | 27,33 | 91,47 | 8,98 | 0.3384 |
| C20:0 | 0,66 | 0,12 | 0,79 | 0,20 | >0.9999 | 0,70 | 0,14 | 0,73 | 0,13 | 0.9787 | 0,74 | 0,33 | 0,76 | 0,08 | 0.9671 |
| C20:1 | 1,03 | 0,14 | 1,17 | 0,56 | >0.9999 | 1,48 | 0,42 | 1,52 | 0,32 | 0.9978 | 1,76 | 0,47 | 1,46 | 0,50 | 0.9287 |
| C20:3 n-6 | 5,68 | 0,82 | 6,10 | 1,70 | 0.9998 | 4,93 | 1,28 | 5,96 | 0,46 | 0.3626 | 6,36 | 1,82 | 8,18 | 4,59 | 0.7188 |
| C20:4 n-6 | 57,73 | 8,22 | 57,46 | 6,90 | 0.9999 | 51,26 | 6,01 | 48,65 | 10,80 | 0.9506 | 61,41 | 19,06 | 74,17 | 36,28 | 0.9482 |
| C22:5 n-3 | 0,94 | 0,23 | 0,95 | 0,29 | >0.9999 | 0,84 | 0,21 | 0,75 | 0,24 | 0.9437 | 1,23 | 0,59 | 0,80 | 0,09 | 0.2439 |
| C22:6 n-3 | 17,70 | 2,32 | 17,17 | 1,03 | 0.9964 | 15,52 | 0,80 | 13,40 | 1,75 | 0.1383 | 18,52 | 4,02 | 19,70 | 6,20 | 0.9997 |
| FA tot | 337,23 | 13,82 | 393,15 | 113,39 | 0.5298 | 337,41 | 72,47 | 340,88 | 15,47 | 0.9997 | 391,75 | 85,03 | 374,64 | 83,60 | 0.7675 |

**Supplementary Table S3. Plasma total fatty acids in 5xFAD male mice at 2, 5 and 8 months of age.** Data are expressed as mean±SEM; 5xFAD vs NC were compared by two-way ANOVA with with Šídák's multiple comparisons test as appropriate.

| Fatty acid | 2 MO | | | | | 5 MO | | | | | 8 MO | | | | |
|---|---|---|---|---|---|---|---|---|---|---|---|---|---|---|---|
| | NC | | 5xFAD | | p value | NC | | 5xFAD | | p value | NC | | 5xFAD | | p value |
| | Mean | SD | Mean | SD | | Mean | SD | Mean | SD | | Mean | SD | Mean | SD | |
| C14:0 | 0,39 | 0,08 | 0,44 | 0,19 | >0.9999 | 0,37 | 0,13 | 0,42 | 0,07 | 0.9874 | 0,54 | 0,23 | 0,53 | 0,13 | >0.9999 |
| C16:0 | 39,89 | 4,57 | 40,24 | 7,57 | >0.9999 | 26,24 | 2,90 | 37,92 | 10,99 | 0.6563 | 36,71 | 5,75 | 36,91 | 4,88 | >0.9999 |
| C16:1 t | 0,80 | 0,15 | 0,92 | 0,20 | >0.9999 | 0,61 | 0,15 | 0,92 | 0,36 | 0.8824 | 1,08 | 0,55 | 0,73 | 0,13 | 0.8667 |
| C16:1 | 2,41 | 0,26 | 2,23 | 0,93 | >0.9999 | 1,33 | 0,04 | 3,67 | 1,70 | 0.4894 | 2,92 | 0,84 | 2,36 | 0,79 | 0.9792 |
| C18:0 | 36,28 | 3,89 | 33,95 | 1,97 | 0.9828 | 27,66 | 0,68 | 30,39 | 2,26 | 0.8225 | 33,63 | 2,83 | 31,34 | 4,06 | 0.8771 |
| C18:1 n-9 | 33,98 | 4,83 | 31,51 | 7,20 | 0.9796 | 18,07 | 4,02 | 38,30 | 19,50 | 0.5168 | 35,62 | 13,82 | 26,56 | 6,97 | 0.8552 |
| C18:1 n-7 | 4,65 | 1,07 | 4,75 | 1,30 | >0.9999 | 3,09 | 0,21 | 6,32 | 2,75 | 0.3789 | 6,07 | 1,62 | 4,26 | 1,22 | 0.7190 |
| C18:2 n-6 | 65,68 | 11,39 | 65,13 | 11,43 | 0.9998 | 41,10 | 15,40 | 64,62 | 24,51 | 0.5085 | 65,29 | 13,96 | 59,34 | 7,54 | 0.9502 |
| C20:0 | 0,25 | 0,04 | 0,29 | 0,08 | >0.9999 | 0,10 | 0,13 | 0,28 | 0,08 | 0.2695 | 0,25 | 0,05 | 0,21 | 0,12 | 0.9814 |
| C20:1 | 0,67 | 0,05 | 0,78 | 0,25 | >0.9999 | 0,40 | 0,08 | 0,92 | 0,43 | 0.4128 | 0,93 | 0,37 | 0,69 | 0,36 | 0.8794 |
| C20:3 n-6 | 3,17 | 0,69 | 2,66 | 0,16 | 0.9998 | 1,72 | 0,17 | 2,68 | 0,75 | 0.5468 | 2,79 | 0,50 | 2,34 | 0,31 | 0.9802 |
| C20:4 n-6 | 40,89 | 4,48 | 42,90 | 3,49 | 0.9888 | 30,83 | 3,63 | 30,48 | 4,01 | >0.9999 | 31,48 | 2,61 | 32,19 | 10,23 | 0.9999 |
| C22:5 n-3 | 0,50 | 0,07 | 0,47 | 0,09 | >0.9999 | 0,20 | 0,28 | 0,47 | 0,19 | 0.4702 | 0,54 | 0,11 | 0,60 | 0,08 | 0.9891 |
| C22:6 n-3 | 10,00 | 0,75 | 11,36 | 1,38 | >0.9999 | 7,48 | 1,80 | 7,92 | 0,87 | 0.9748 | 9,06 | 1,22 | 8,78 | 2,15 | 0.9989 |
| FA tot | 244,25 | 17,84 | 241,66 | 26,22 | 0.9767 | 161,41 | 20,21 | 230,65 | 57,90 | 0.4291 | 231,56 | 37,24 | 210,53 | 22,20 | 0.9392 |

**Supplementary Table S4. Plasma total fatty acids in 5xFAD female mice at 2, 5 and 8 months of age.** Data are expressed as mean±SEM; 5xFAD vs NC were compared by two-way ANOVA with with Šídák's multiple comparisons test as appropriate.

| Fatty acid | NC | | 5xFAD | | p value | NC | | 5xFAD | | p value |
|---|---|---|---|---|---|---|---|---|---|---|
| | Mean | SD | Mean | SD | | Mean | SD | Mean | SD | |
| C14:0 | 0,07 | 0,02 | 0,08 | 0,02 | 0.9808 | 0,11 | 0,05 | 0,09 | 0,02 | 0.6227 |
| C16:0 | 10,24 | 0,63 | 10,88 | 0,59 | 0.2658 | 10,16 | 0,45 | 10,57 | 1,09 | 0.5712 |
| C16:1 t | 0,07 | 0,01 | 0,07 | 0,02 | 0.9868 | 0,07 | - | 0,08 | 0,01 | 0.0639 |
| C16:1 | 0,26 | 0,03 | 0,29 | 0,02 | 0.1952 | 0,25 | 0,03 | 0,28 | 0,04 | 0.1392 |
| C18:0 | 8,64 | 0,37 | 9,10 | 0,25 | 0.2008 | 8,85 | 0,47 | 8,87 | 0,69 | 0.9981 |
| C18:1 n-9 | 9,71 | 1,46 | 9,91 | 0,69 | 0.9421 | 9,38 | 0,63 | 9,82 | 1,39 | 0.7462 |
| C18:1 n-7 | 1,58 | 0,12 | 1,76 | 0,07 | 0.9166 | 1,63 | 0,14 | 1,71 | 0,17 | 0.5789 |
| C18:2 t | 0,71 | 0,12 | 0,73 | 0,06 | 0.9181 | 0,74 | 0,08 | 0,74 | 0,10 | 0.9960 |
| C18:2 n-6 | 0,32 | 0,06 | 0,32 | 0,02 | 0.9921 | 0,32 | 0,04 | 0,32 | 0,05 | 0.9880 |
| C20:0 | 0,08 | 0,01 | 0,09 | 0,01 | 0.5331 | 0,08 | 0,01 | 0,09 | 0,01 | 0.5447 |
| C20:1 | 0,75 | 0,14 | 0,75 | 0,04 | 0.9996 | 0,78 | 0,11 | 0,76 | 0,11 | 0.9217 |
| C20:3 n-6 | 0,19 | 0,02 | 0,21 | 0,02 | 0.0965 | 0,18 | 0,02 | 0,17 | 0,01 | 0.7469 |
| C20:4 n-6 | 3,88 | 0,20 | 4,19 | 0,20 | 0.0925 | 4,02 | 0,31 | 4,19 | 0,31 | 0.4613 |
| C22:3 | 0,88 | 0,04 | 0,88 | 0,06 | 0.9936 | 0,98 | 0,10 | 0,89 | 0,07 | 0.1047 |
| C22:5 n-3 | - | - | 0,05 | 0,04 | 0.0237 | 0,02 | 0,03 | 0,01 | 0,03 | 0.9217 |
| C22:6 n-3 | 7,29 | 0,12 | 7,65 | 0,66 | 0.4689 | 7,38 | 0,54 | 7,25 | 0,70 | 0.9074 |
| FA total | 45,90 | 3,05 | 48,13 | 2,43 | 0.4515 | 46,01 | 2,61 | 47,10 | 4,71596104 | 0.8203 |

**Supplementary Table S5. Cortex total fatty acids in 5xFAD male and female mice at 8 months of age.** Data are expressed as mean±SEM; 5xFAD vs NC were compared by two-way ANOVA with with Šídák’s multiple comparisons test as appropriate.

| Fatty acid | NC-Vehicle | | 5xFAD-Vehicle | | NC-SCDi | | 5xFAD-SCDi | | P value |
|---|---|---|---|---|---|---|---|---|---|
| | Mean | SD | Mean | SD | Mean | SD | Mean | SD | |
| C14:0 | 0,08 | 0,01 | 0,10 | 0,01 | 0,07 | 0,01 | 0,08 | 0,01 | NC-V vs 5xFAD-V p=0.0212; NC-S vs 5xFAD-V p=0.0014 |
| C16:0 | 10,47 | 1,00 | 10,97 | 0,70 | 9,99 | 0,31 | 11,56 | 0,31 | NC-V vs 5xFAD-S p=0.0343; NC-S vs 5xFAD-S p=0.005 |
| C16:1 t | 0,08 | 0,02 | 0,11 | 0,01 | 0,08 | 0,01 | 0,11 | 0,01 | NC-V vs 5xFAD-V p=0.0026; NC-V vs 5xFAD-S p=0.0045; NC-S vs 5xFAD-V p=0.0008; NC-S vs 5xFAD-S p=0.0013 |
| C16:1 | 0,29 | 0,05 | 0,36 | 0,06 | 0,27 | 0,02 | 0,36 | 0,02 | NC-V vs 5xFAD-S p=0.0051; NC-S vs 5xFAD-V p=0.0240; NC-S vs 5xFAD-S p=0.0065 |
| C18:0 | 9,61 | 0,82 | 9,34 | 1,29 | 9,42 | 0,40 | 10,37 | 0,37 | - |
| C18:1 n-9 | 12,81 | 2,20 | 14,23 | 2,03 | 12,70 | 1,50 | 13,88 | 0,67 | - |
| C18:1 n-7 | 1,59 | 0,19 | 1,79 | 0,14 | 1,55 | 0,06 | 1,85 | 0,06 | NC-V vs 5xFAD-S p=0.0143; NC-S vs 5xFAD-S p=0.0067; NC-S vs 5xFAD-V p=0.0014; NC-S vs 5xFAD-S p=0.0018 |
| C18:2 t | 0,26 | 0,06 | 0,29 | 0,12 | 0,19 | 0,05 | 0,20 | 0,02 | - |
| C18:2 n-6 | 0,46 | 0,09 | 0,48 | 0,09 | 0,44 | 0,07 | 0,47 | 0,02 | - |
| C20:0 | 0,14 | 0,03 | 0,15 | 0,06 | 0,14 | 0,02 | 0,15 | - | - |
| C20:1 | 1,01 | 0,19 | 1,11 | 0,31 | 0,98 | 0,10 | 1,10 | 0,07 | - |
| C20:3 n-6 | 0,17 | 0,06 | 0,20 | 0,01 | 0,16 | 0,01 | 0,22 | 0,02 | NC-V vs 5xFAD-V p=0.0085; NC-V vs 5xFAD-S p=0.0003; NC-S vs 5xFAD-V p=0.0015; NC-S vs 5xFAD-S p<0.0001 |
| C20:4 n-6 | 4,89 | 0,39 | 5,03 | 0,46 | 4,88 | 0,20 | 5,47 | 0,38 | NC-V vs 5xFAD-S p=0.0465; NC-S vs 5xFAD-S p=0.0417 |
| C22:3 | 1,14 | 0,05 | 1,21 | 0,05 | 1,15 | 0,05 | 1,29 | 0,13 | NC-V vs 5xFAD-S p=0.0158; NC-S vs 5xFAD-S p=0.0262 |
| C22:5 n-3 | 0,06 | - | 0,08 | 0,01 | 0,06 | - | 0,09 | - | NC-V vs 5xFAD-V p<0.0001; NC-V vs 5xFAD-S p<0.0001; NC-S vs 5xFAD-V p<0.0001; 5xFAD-V vs 5xFAD-S p=0.0435; NC-S vs 5xFAD-S p<0.0001 |
| C22:6 n-3 | 6,24 | 0,54 | 6,29 | 0,39 | 6,22 | 0,47 | 6,97 | 0,17 | NC-V vs 5xFAD-S p=0.0298; 5xFAD-V vs 5xFAD-S p=0.039; NC-S vs 5xFAD-S p=0.0257 |
| FA tot | 51,38 | 5,75 | 54,07 | 4,39 | 50,39 | 2,52 | 56,36 | 1,85 | - |

**Supplementary Table S6. Hippocampal total fatty acids in 5xFAD SCDi treated mice.** Data are expressed as mean±SEM; 5xFAD vs NC were compared by two-way ANOVA with Fisher's LSD post-hoc test as appropriate. Only significant p values (p<0.01) are shown, - displays no significant difference between groups.

| Lipid classes | Internal standard | Supplier | Cat. Number | Concentration (mg/mL) |
|---|---|---|---|---|
| CE | CE 17:0 (Cholesteryl heptadecanoate >99%) | Nu-Chek- Prep, Inc | CH-816-S21-B | 0.02 |
| TG | TG 19:0 (Trinonadecanoin >99%) | Nu-Chek- Prep, Inc | T-165-J6-C | 0.02 |
| FFA | FFA 24:0 (Lignoceric acid >99%) | Sigma-Aldrich | L6641 | 0.094 |
| PL | PC 15:0 (1,2-dipentadecanoyl-sn-glycero-3-phosphocholine >99%) | Avanti Polar Lipids, Inc | 850350P | 0.06 |

**Supplementary Table S7. GC-FID internal standards.** Internal standards used for each lipid class and respective concentrations.
